# Predicting the functional effect of compound heterozygous genotypes from large scale variant effect maps

**DOI:** 10.1101/2023.01.11.523651

**Authors:** Michael J. Xie, Gareth A. Cromie, Katherine Owens, Martin S. Timour, Michelle Tang, J. Nathan Kutz, Ayman W. El-Hattab, Richard N. McLaughlin, Aimée M. Dudley

## Abstract

**Background:** Pathogenic variants in *PHGDH, PSAT1*, and *PSPH* cause a set of rare, autosomal recessive diseases known as serine biosynthesis defects. Serine biosynthesis defects present in a broad phenotypic spectrum that includes, at the severe end, Neu–Laxova syndrome, a lethal multiple congenital anomaly disease, intermediately in the form of infantile serine biosynthesis defects with severe neurological manifestations and growth deficiency, and at the mild end, as childhood disease with intellectual disability. However, because L-serine supplementation, especially if started early, can ameliorate and in some cases even prevent symptoms, knowledge of pathogenic variants is highly actionable.

**Methods:** Recently, our laboratory established a yeast-based assay for human *PSAT1* function. We have now applied it at scale to assay the functional impact of 1,914 SNV-accessible amino acid substitutions. In addition to assaying the functional impact of individual variants in yeast haploid cells, we can assay pairwise combinations of *PSAT1* alleles that recapitulate human genotypes, including compound heterozygotes, in yeast diploids.

**Results:** Results of our assays of individual variants (in haploid yeast cells) agree well with clinical interpretations and protein structure-function relationships, supporting the use of our data as functional evidence under the ACMG interpretation guidelines. Results from our diploid assay successfully distinguish patient genotypes from those of healthy carriers and agree well with disease severity. Finally, we present a linear model that uses individual allele measurements (in haploid yeast cells) to accurately predict the biallelic function (in diploid yeast cells) of ^~^1.8 million allele combinations corresponding to potential human genotypes.

**Conclusions:** Taken together, our work provides an example of how large-scale functional assays in model systems can be powerfully applied to the study of a rare disease.

## Background

High throughput sequencing technologies have accelerated the discovery of human genetic variation. Data from the Genome Aggregation Database (gnomAD) [1] suggest that the average person harbors approximately 200 rare (allele frequency <0.1%) protein coding variants [2]. However, leveraging genomic information for the diagnosis and treatment of medically actionable diseases is currently limited by the relatively small number of alleles for which clinical interpretation is readily available. Most of the ^~^550K missense variants in ClinVar [3], even those in well-studied disease genes, are variants of uncertain significance (VUS). The difficulty of variant interpretation is compounded further in rare diseases, and patients often face long diagnostic odysseys or remain undiagnosed.

Inborn errors of metabolism (IEM), genetic alterations that disrupt metabolic homeostasis, are a relatively large class of rare diseases. Over 700 IEMs [4] have been described and new disorders continue to be discovered [5]. Although individually rare, collectively these diseases are common, with estimates suggesting that IEMs may affect 1 in 800-2600 live births [6,7]. Many are severe and manifest early in life, but some are also medically actionable and timely diagnosis can prevent the onset of irreversible damage. In these diseases, disruptions of metabolic pathways can lead to toxic levels of substrate accumulation or deficiency of an essential product. Therapeutic strategies often focus on amending these imbalances. For example, dietary restriction is prescribed for the treatment of urea cycle disorders, phenylketonuria, and galactosemia, while dietary supplementation is prescribed for homocystinuria and pyridoxine-dependent epilepsy [8–10].

Serine biosynthesis defects are a group of clinically actionable IEMs of varying severity that were first described in the 1970’s, characterized biochemically in the 1990’s, and mapped genetically in the 2000’s (**Fig. 1**). Impairment of any of the three L-serine biosynthesis pathway enzymes, phosphoglycerate dehydrogenase (PGDH; encoded by *PHGDH*), phosphoserine aminotransferase (PSAT; encoded by *PSAT1*), and phosphoserine phosphatase (PSP; encoded by *PSPH*), results in systemic serine deficiency [11]. Serine metabolism is central to numerous biological processes, including the synthesis of proteins, nucleotides, and phospholipids, as well as the formation of the neuromodulators D-serine and glycine. As a result, serine biosynthesis defects negatively impact nervous system development and function, with clinical manifestations that include microcephaly, seizures, intellectual disability, and neuropathies. Case studies have demonstrated that oral serine supplementation can reduce and, in some cases, prevent the onset of these severe symptoms that typically manifest very early in life [12–16]. Prompt diagnosis is crucial, as the impact of therapeutic intervention, including prenatal dietary supplementation, is greatest before patients become symptomatic and irreversible neurological damage occurs.

**Fig. 1.**
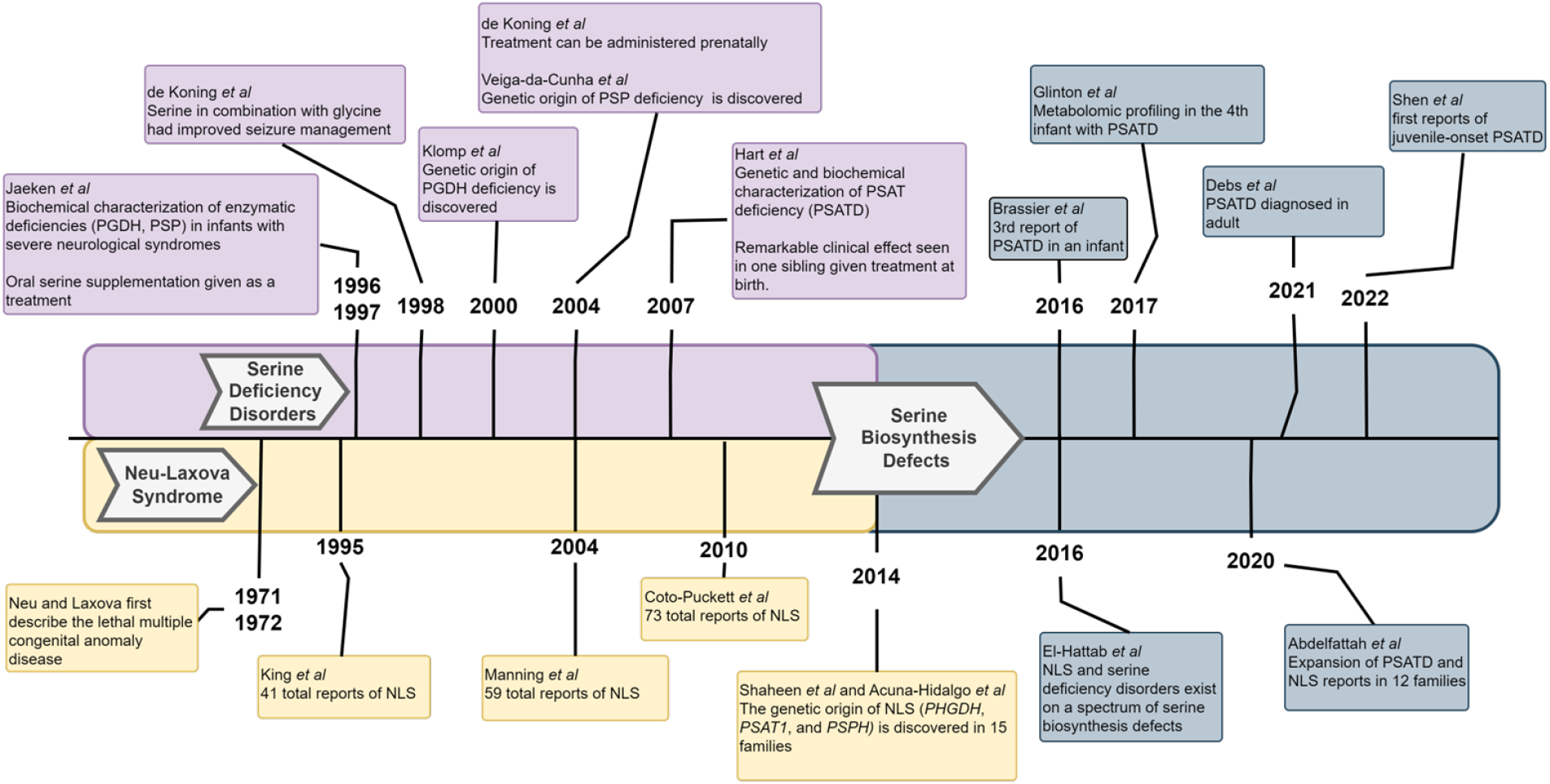
Key developments in the study of serine deficiency disorders, a set rare but treatable inborn errors of metabolism [11–30].

Unfortunately, like many rare diseases, diagnosing serine deficiency disorders is difficult in the absence of a family history of disease. One challenge is that the disorders display a broad phenotypic spectrum (reviewed in [11,31]). At one extreme, Neu-Laxova syndrome (NLS) is a severe, lethal, multiple congenital anomaly disease characterized by marked intrauterine growth restriction, microcephaly, distinctive facial features, limb and skin defects, and variable brain malformations including lissencephaly, corpus callosum agenesis, and hypoplastic cerebellum and pons. Infantile serine biosynthesis defects represent the intermediate form of the disorder spectrum, typically presenting with intrauterine growth restriction, microcephaly, feeding difficulties, vomiting, irritability, hypertonia evolving into spastic tetraplegia, nystagmus, early onset seizures, and poor psychomotor development. Other findings include congenital cataracts, adducted thumbs, megaloblastic anemia, brain atrophy, hypomyelination, and hypoplastic cerebellum and pons. At the mild end of the disorder spectrum are childhood serine biosynthesis defects characterized by developmental delay, intellectual disability, and behavioral abnormalities. Other findings include epilepsy, ataxia, nystagmus, polyneuropathy, and hypertonia. Although it has been hypothesized that residual enzyme activity may explain the phenotypic variability [25,28,32] within serine deficiency disorders, functional studies regarding this subject are sparse and the natural histories of the disorders are largely unknown. Biochemical diagnosis of serine biosynthesis defects can be challenging as it requires measurement of amino acid levels in patient serum or cerebral spinal fluid (CSF) with comparison to closely age-matched controls. Although the serum-directed metabolic screen is more widely used, considered generally in children with intellectual disability, serine and glycine plasma concentrations can be normal in the presence of serine biosynthesis defects if the blood sample is not obtained in a fasted state. In contrast, CSF serine and glycine concentrations are not affected by meals, however, CSF amino acid analysis is not typically done in the absence of seizures. Therefore, the metabolic work up may fail to identify children with serine biosynthesis defects [27,33].

Identifying the presence of pathogenic sequence variants through whole exome/genome sequencing has several potential advantages for diagnosing inherited metabolic diseases, such as serine biosynthesis defects. However, this approach relies on having clinical interpretations for rare variants, which are generally not available. Instead, within the current disease literature, clinical sequencing has primarily been used as a tool by experts to provide molecular confirmation once a patient has become symptomatic. High-throughput approaches to variant interpretation that can be applied to rare variants are needed to increase the success rate of sequencing-based diagnostics. Although computational prediction methods can be applied at scale, their relatively high error rates limit their utility [34]. Functional assays that can quantitatively measure variant effects on protein activity [35] are an alternative approach that can be used as part of the criteria for variant interpretation established by the American College of Medical Genetics (ACMG) [36]. Improvements in DNA synthesis technology have enabled time-and-cost effective methods for building large variant libraries. When combined with high-throughput phenotyping methods, functional assays can comprehensively assess variants that have been identified in the human population as well as those that may arise in the future [37,38].

Recently, our laboratory established a yeast-based assay for human PSAT function [39]. PSAT catalyzes the reversible conversion of glutamate to α-ketoglutarate and 3-phosphohydroxypyruvate (3-PHP) to phosphoserine [40]. PSAT belongs to a large family of fold type I pyridoxal phosphate (PLP; the active form of vitamin b6) cofactor-dependent enzymes, and more specifically, the class IV group of aminotransferases [41,42]. The crystal structure of human PSAT complexed with PLP (PDB: 3e77; residues L17-L370) shows that the protein adopts an s-shaped homodimer assembly in which two separate PLP-containing active sites surrounded by an overall positive charge distribution reside along the interface (Fig. 2B). Each subunit of the homodimer consists of two domains; a large domain, which forms the dimer interface and binds PLP, and a small domain (Fig. 2A). Despite extensive literature on the structure and biochemistry of PSAT [40], there is not a comprehensive understanding of which mutations at which positions impair the ability of PSAT to support normal serine biosynthesis.

**Fig. 2.**
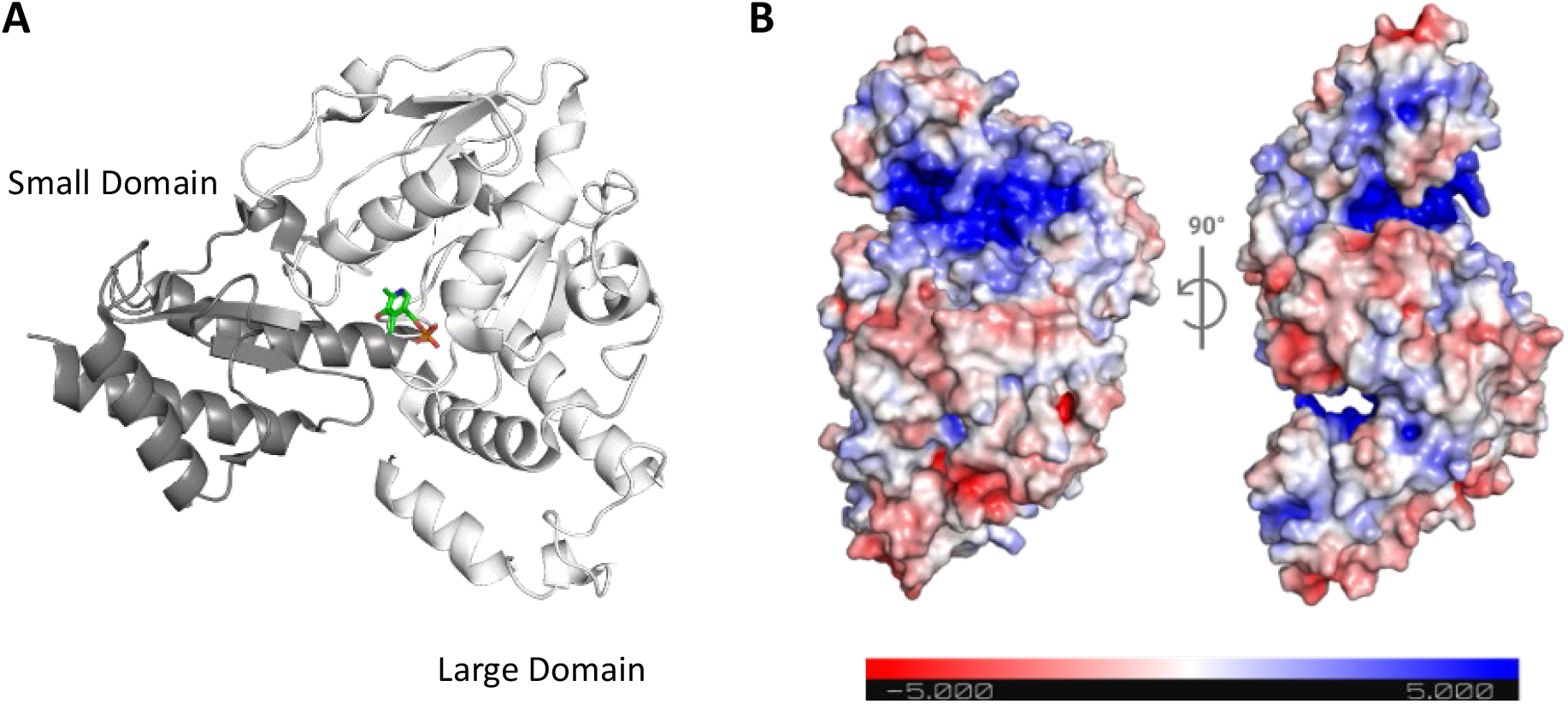
Overall subunit organization and homodimer assembly of human PSAT. **A** Ribbon diagram representation of the secondary structure of a subunit of PSAT (PDB 3e77; L17-L370) complexed with PLP (green) in stick representation. The small C-terminal domain is colored as dark grey and large N-terminal domain as white. **B** Local charge distribution on the surface of the PSAT homodimer assembly, on a scale of negative charge (−5 kT; red) to positive charge (+5 kT; blue) as calculated by the Adaptive Poisson–Boltzmann Solver program.

Here, we apply our yeast-based assay at scale to quantify the functional impact of 1,914 amino acid substitutions that are accessible via a single nucleotide variant (SNV) in the human *PSAT1* coding sequence. Our results agree well with clinically interpreted alleles and with protein structure-function relationships, supporting the use of our data as functional evidence under the ACMG interpretation guidelines. In addition to assaying the functional impact of single variants in yeast haploids, we construct and assay yeast diploids with pairwise combinations of *PSAT1* alleles that recapitulate human genotypes. The results of this diploid functional assay distinguish patient genotypes from those of healthy carriers and agree with the stratification of patient genotypes by disease severity. Finally, we develop a mathematical model that can accurately predict biallelic function in diploids from pairwise combinations of individual allele activity measured experimentally in haploids. Taken together, our work provides an example of how large-scale functional assays in model systems can be powerfully applied in the study of rare disease and to inform future diagnostic efforts.

## Methods

### Strain library construction

All *Saccharomyces cerevisiae* strains used in this study (Additional File 1: Tables S1 and S2) were derived from the isogenic lab strains FY4 (MAT**a**) and FY5 (MATα) [43]. Unless otherwise noted, strains were grown in rich medium (YPD, 1% yeast extract, 2% peptone, and 2% glucose) or minimal medium (SD, without amino acids, 2% glucose) using standard media conditions and methods for yeast genetic manipulation [44].

Methodology concerning the design and construction of the yeast codon-optimized version (*yPSAT1*) of human *PSAT1* isoform 1, the wild type (*yPSAT1*) strain, the deletion (*ser1Δ0*) strain, and individual variant strains were previously described in Sirr *et al* [39]. Here, we describe the design and construction of a variant library ^~^10 fold larger and modifications of the growth assay for high throughput as follows:

The variant library was designed to capture the amino acid substitutions resulting from all SNV-accessible missense mutations across 369 codons, excluding the start and stop, in the human *PSAT1* isoform 1 cDNA sequence (CCDS6660.1; Consensus Coding Sequence database [45]). The 9 possible single nucleotide variants at each codon resulted in 4-7 unique amino acid substitutions. *yPSAT1* variants encoding the complete set of these unique amino acid substitutions (n = 2,182) were synthesized (Twist Biosciences) in *yPSAT1* as an oligonucleotide library in which each well of a 96 well plate contained all amino acid substitutions (n=4-7) at a given amino acid.

Individual wells of these plates were amplified to approximately 500 ng of DNA by 15-cycles of high-fidelity PCR (Phusion High-Fidelity DNA Polymerase; Thermo Scientific) and transformed into a MAT**a** haploid deletion strain (*ser1Δ0*) using standard methods. Single colonies from the yeast transformations were isolated such that 6,335 individual transformants, each encoding a single amino acid substitution, were arrayed into 96-well plates containing rich medium. Because individual transformants are isolated and maintained as separate stocks (one strain per well in a 96-well plate), each strain is an independently constructed biological isolate of the variant it contains. For downstream phenotype normalization, each library plate also contained replicates of the same control strains: 2 deletion (*ser1Δ0*) and 4 wild type (*yPSAT1*) strains.

### Variant library sequence confirmation

Because of the transformation approach described above, for each *yPSAT1* transformant, we know which codon is mutated (target codon), but not which specific variant is present. To determine this, we used a custom MinION (Oxford Nanopore Technologies) sequencing pipeline (described in detail in Additional File 2). Briefly, individual transformants were pooled in groups of 12, so that no target codon was represented more than once in a single pool. Each pool was then sequenced and, at each target codon, the most frequent potential variant codon was identified (candidate variant) as well as the second most frequent variant. Because we know which target codon corresponds to which transformant, this allows us to associate each candidate variant with a single transformant.

For any given DNA sequence, the frequency and pattern of MinIon sequencing errors varies greatly from base to base. These errors can occur at frequencies high enough to generate spurious matches to variant codons. However, the error patterns are also reproducible, allowing us to develop an error model for each variant, describing the frequency with which it is generated by sequencing errors. This frequency can be compared to the frequency observed for each candidate variant in the pooled sequencing, allowing true variants to be distinguished from sequencing noise.

Using this approach, candidate variants underwent quality control. Candidates were rejected if the observed frequency of the variant was less than 3.3× the estimated error frequency for that variant. In addition, candidates were also rejected if the second most frequent variant was both enriched (>=10-fold) relative to its error frequency and was observed at >30% of the frequency of the candidate variant. Finally, candidates were rejected if they were supported less than 15 reads, or if any missense or nonsense secondary mutations were present in the *yPSAT1* sequence. The end result of this process was that each transformant was assigned either a high confidence call for the variant present in that transformant, or an NA call that resulted in that transformant being removed from analysis.

### Validating the MinION Variant Calls Using Illumina Sequencing

To validate our Oxford Nanopore pipeline, we performed amplicon sequencing on the Illumina sequencing platform to confirm the variant in each well of each variant plate. These calls were then compared to the result from the Oxford Nanopore pipeline. For the Illumina sequencing, the *yPSAT1* ORF was divided up into five ^~^300bp overlapping segments. The specific segment to be amplified and sequenced corresponded to the known codon position of the variant within our library. 35 cycles of PCR amplification were performed using genomic DNA as a template. The initial PCR reaction was then diluted and used for an additional 24 cycles of PCR to add on Illumina index sequences. The final PCR reactions were pooled and gel purified before performing Illumina sequencing on a Nextseq 500 using 300-cycle, dual-indexed, paired-end sequencing. Reads were demultiplexed and aligned to the *yPSAT1* reference sequence as described [39]. For each isolate, the basecalls across the target codon position were used to identify the variant codon sequence.

Comparison of the MinION and Illumina results revealed excellent agreement (95.5%) for the MinION calls passing the quality filters (Additional File 2: Fig. S1). Most of the “variants” failing the MinION quality filters are occasions when, as determined by the Illumina sequencing, no change had actually been made at the target codon (i.e. wild type *yPSAT1* sequence). As expected, for these “no-change” isolates, the most frequent potential variant identified by the MinION pipeline occurred at a low frequency similar to the error frequency for that variant (Additional File 2: Fig. S1A). In contrast, when a variant was present, it was observed at higher frequency and was correctly identified with high accuracy.

The accession number for the Illumina and MinION read sequences generated for this study is (in process, available upon manuscript acceptance).

### Generation of Yeast Diploids

Unless noted, all *yPSAT1* variants are in a single mating type (MAT**a**) (Additional File 1: Table S1). To generate diploid strains harboring combinations of *yPSAT1* alleles (missense variant, wild type, or null), variants were first introduced into a strain of the opposite mating type. The resulting *yPSAT1* MATα strains were then mated in a pairwise manner to the relevant *yPSAT1* MAT**a** strains using standard methods [46]. The resulting diploid strains, harboring combinations of *yPSAT1* alleles (Additional File 1: Table S2), were then arrayed in an alternating checkerboard pattern that minimizes the influence of nutrient competition from neighboring colonies during phenotype measurement.

### Growth assays

Before phenotyping, the set of variant strains generated in this study were extended to include the haploid strains carrying missense variants constructed and sequence confirmed by Sirr *et al* [39]. These additional strains were re-arrayed to match the phenotyping layout used in this study, with at least 2 isolates of each variant. Additional control plates included a plate with wild type at every position, and plates consisting entirely of D100A or A99V (with control positions) *yPSAT1* strains, which are ClinVar pathogenic alleles encoding missense variants.

A Biomek i7 robot outfitted with a V&P 96-pin head was utilized to pin strain plate libraries between different culture media. Strains were initially grown to saturation in rich medium and then pinned onto solid medium utilizing glycerol as the central carbon source (YPG,1% yeast extract, 2% peptone, 2% glycerol) to remove any yeast cells lacking mitochondria (petite). Strains were then pinned back into rich medium and grown to saturation. Each plate was then pinned in replicate (n=3-6) onto solid minimal medium, which lacks serine, and grown for 3 days at 30°C. A mounted Canon PowerShot SX10 IS compact digital camera was used to take images (ISO200, f4.5, 1/40s exposure) every 24 hours for three days under consistent lighting, camera to subject distance, and zoom. Each plate was labeled with a custom code39 barcode that was included in the frame of view. Images were acquired as jpg files.

### Image-based growth quantification

Each plate image was processed using PyPl8 (https://github.com/lacyk3/PyPl8) to extract features from each strain patch in each barcoded agar plate as follows: First, the barcode within each image was detected and decoded to rename each file using the corresponding plate name, replicate number, condition, and timepoint. Next, each image was cropped into 96 square tiles, segmented, and each replica pinned patch was identified using Otsu’s method or circle detection. Finally, the sum of the gray scale pixel intensities within each strain patch (pixelsum), was extracted and used as the metric for growth estimation.

### Growth data fitting and normalization

For haploid strains, raw phenotypic values were normalized, quality control filters were applied to each isolate, and a final relative growth estimate for each variant was determined, as described in Lo *et al* [47]. Briefly, normalization steps were carried out to account for the effects of plate-to-plate variation, relative growth of neighboring patches, and plate edge effects. Pin effect normalization did not reduce noise and was omitted. Isolates with (nonsynonymous) secondary mutations were removed from the dataset as were all isolates of variants that showed a high degree of variation in isolate-to-isolate growth values. This left a final filtered dataset of 5,164 independent isolates. Finally, a linear model was used to estimate the relative growth of each genotype, on a scale with growth of null controls set to 0 and growth of wild type set to 1. The script carrying out the growth normalization steps is provided as Additional File 4.

A similar approach was applied to phenotypic values extracted from the diploid growth assay, although neighbor effects were assumed negligible because of the checkerboard pinning arrangement. For normalization, a linear model was used to simultaneously estimate the effects of genotype, plate edge positioning and plate-to-plate variation on growth. Genotype effects were rescaled to set homozygous null to 0 and homozygous wild type to 1.

### Predicting diploid growth

We developed a linear model to predict the growth of diploid strains based on which pair of *yPSAT1* variants is present, using individual growth estimates of each allele in haploids. In this model, for each diploid, the (haploid) growth estimates of the lower and higher growth alleles were labelled as minimum and maximum (*j, k*), respectively. In cases of homozygous combinations, the minimum and maximum were equal. Strains carrying a single copy of the wild type allele (*yPSAT1*) or null (*ser1*Δ0) had their respective haploid estimates set equal to 1 and 0. To predict diploid growth (d) from the more impaired (*x_j_*) and less impaired (*x_k_*) alleles, we performed an ordinary least squares regression to fit an additive pairwise model (d = a + *bx_j_* + *cx_k_*), with resulting coefficients being a=0.05, b=0.28, and c =0.73. We also compared this model to two simpler regression models. In the first model, diploid growth was predicted from the mean of the haploid estimates (*x_j_* = *x_k_*). In the second, diploid growth was predicted from the higher growing haploid allele only, i.e. complete dominance (*x_j_* = 0). Leave one out cross validation was used to assess model performance in the best model.

Next, we evaluated the performance of experimental versus predicted growth as a binary classifier for identifying genotypes matching those of patients diagnosed with *PSAT1*-related serine biosynthesis defects vs carrier parents. We fit a logistic regression model to both experimentally measured diploid growth values (Log(p/1-p) = e + f*y_e_*) and predicted diploid growth values (Log(p/1-p) = g + h*y_p_*) calculated from the additive pairwise model. The resulting coefficients for the regression models were e=−7.7, f=10.7, g=−20.0, and h=27.2. The logistic regression models indicated a threshold (p=0.5) decision value of 72% and 73% for experimental and predicted diploid growth, respectively.

### Protein structure and conservation analysis

Structural features were derived from the crystal structure of human phosphoserine aminotransferase isoform 1 complexed with pyridoxal phosphate cofactor (PDB: 3e77; subunit L17-L370, homodimer biological assembly). Secondary structure features were extracted from this crystal structure using DSSP software [48,49]. AlphaFold (version 2022-11-01) [50,51] was used to predict the structure of the missing 16 N-terminal residues. Molecular visualizations were created using the PyMol Molecular Graphics System Version 2.3.2 (Schrödinger). Tools from the publicly available PyMol script and plugin repository were used to determine interfacial residues and charge distribution. Dimer interface residues were defined as residues at which the solvent-exposed surface area in the monomeric model is greater than the solvent-exposed surface area in the dimer model (cutoff value = 0.5 Å^2^). Macromolecular electrostatics were estimated for each residue of PDB 3e77 using the Adaptive Poisson-Boltzmann Solver PyMol plugin. Evolutionary conservation scores and ‘grades’ for each position of the full-length amino acid sequence of human PSAT (UniProtID [52]: Q9Y617-1) were computed using ConSurf [53]. All parameters for the ConSurf calculation were the same as the methodology outlined in creating the ConSurf-DB repository [54], with the best evolutionary model determined to be WAG. The evolutionary conservation scores and ‘grades’ represent the calculated positional conservation based on an amino acid alignment of 300 diverse homologs of PSAT. The determined structure and conservation features for each amino acid position is provided in Additional File 1: Table S6.

## Results

### Surveying the functional impact of large-scale missense variation in *PSAT1*

Our laboratory had previously established a yeast-based complementation assay that leveraged the ability of the human *PSAT1* coding sequence to functionally replace its yeast ortholog, *SER1* [39]. In this assay, growth on minimal medium lacking serine provides a quantitative readout of PSAT activity, allowing the functional impact of protein coding variants to be assessed. Variant impact is expressed on a relatively intuitive scale of activity between the level of yeast growth associated with no activity (that of a complete gene deletion) and that conferred by the wild type human protein coding sequence (*yPSAT1*). Variants causing a reduction in PSAT activity could have their effect via decreases in enzymatic activity, protein stability, or a combination of both.

In this study, we applied the assay at scale to measure the effect of thousands of amino acid substitutions as follows. First, with the exception of the translational initiation and termination codons, we identified all unique amino acid substitutions (n = 2,182) that were accessible via a SNV across the full length of human *PSAT1* isoform 1 cDNA (1,113 bp). Variant codons encoding each amino acid substitution were then introduced into the yeast codon-optimized version of *PSAT1* (*yPSAT1*). The resulting variant library was transformed into a haploid *SER1* deletion strain (*ser1Δ0*) and integrated in single copy at the *SER1* locus of the yeast genome, under the control of the endogenous *SER1* transcriptional promoter and terminator. Next, transformants were individually arrayed in 96-well plate format, and the identity of the variant codon present in each transformant was determined by Oxford Nanopore MinION sequencing (Methods). Strains that harbored secondary mutations in the protein coding sequence were removed from further consideration. The arrayed strain library was then grown, in triplicate, on solid medium lacking serine and imaged after three days at 30°C. Data from these images were used to measure the growth of each strain, relative to wild type (*yPSAT1*) and null (*ser1Δ0*), using a custom automated image analysis pipeline and normalization software (Methods).

Here, we report the functional impact of 1,914 amino acid substitutions in *PSAT1* (Additional File 1: Table S3), corresponding to ^~^88% of all unique SNV-accessible amino acid substitutions (n=2,182) in the human *PSAT1* cDNA sequence. A subset of these substitutions (n=196) was also assayed in our previous study [39] which included the full set of *PSAT1* missense variants described in ClinVar, gnomAD, or the clinical literature at that time. Reanalysis in the current study allowed us to assess these amino acid substitutions using an improved data processing pipeline as the 1,718 new substitutions, thereby placing them on a common scale to facilitate direct comparison. Despite the use of slightly different data analysis pipelines, there was excellent agreement (R^2^=0.94) between the normalized growth estimates of the 196 substitutions in the two studies.

Among the full set of 1,914 amino acid substitutions, the distribution of growth values was bimodal, with a large peak centered near the value of the wild type *yPSAT1* strain (normalized growth=1) and a smaller peak centered around the value of the null (deletion) strain (normalized growth=0) (Fig. 3). These results are consistent with a large group of protein coding variants showing little or no functional impairment relative to wild type, and a smaller group behaving as complete loss of function alleles, comparable to the null control. The remaining protein coding variants showed varying levels of functional impairment (Fig. 3). On this basis, we classified amino acid substitutions in our assay as follows. Substitutions with ≤95% normalized growth were considered functionally impaired relative to wild type and the remaining substitutions (>95% normalized growth) were considered unimpaired. Among the impaired class, we further defined any substitutions resulting in ≤5% normalized growth (i.e. comparable to that of the null control) as amorphs, and any substitutions resulting in less severe functional impairment (>5% and ≤95% normalized growth) as hypomorphs.

**Fig. 3.**
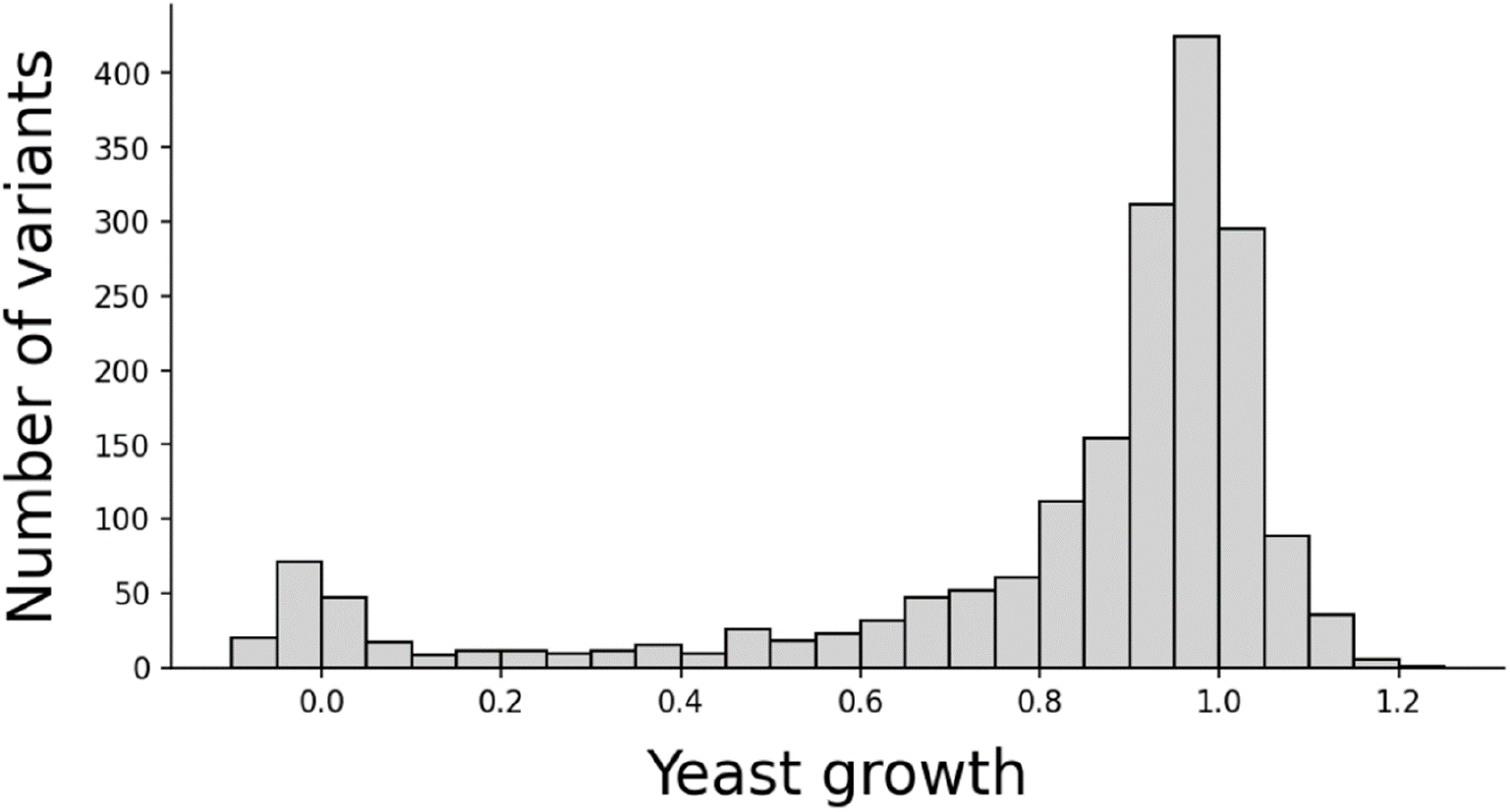
Distribution of variant effect for 1,914 PSAT1 unique SNV-accessible missense amino acid substitutions. Frequency (in 5% intervals) of experimentally measured yeast growth values scaled relative to wild type yPSAT1 (normalized growth=1) and null (normalized growth=0).

### Mapping functional effects to protein structure and conservation

Our dataset of 1,914 PSAT variants assesses the functional impact of the 4-7 SNV-accessible amino acid substitutions at each position across the length of the human PSAT protein (370 aa). At some positions, the majority of SNV-accessible variants exhibited some degree of functional impairment, while other positions were mutationally tolerant to all sampled variants (Fig. 4A). To provide a better understanding of the impact of missense substitutions on protein function, we examined the results of our assay in the context of evolutionary and structural features of the protein [40].

**Fig. 4.**
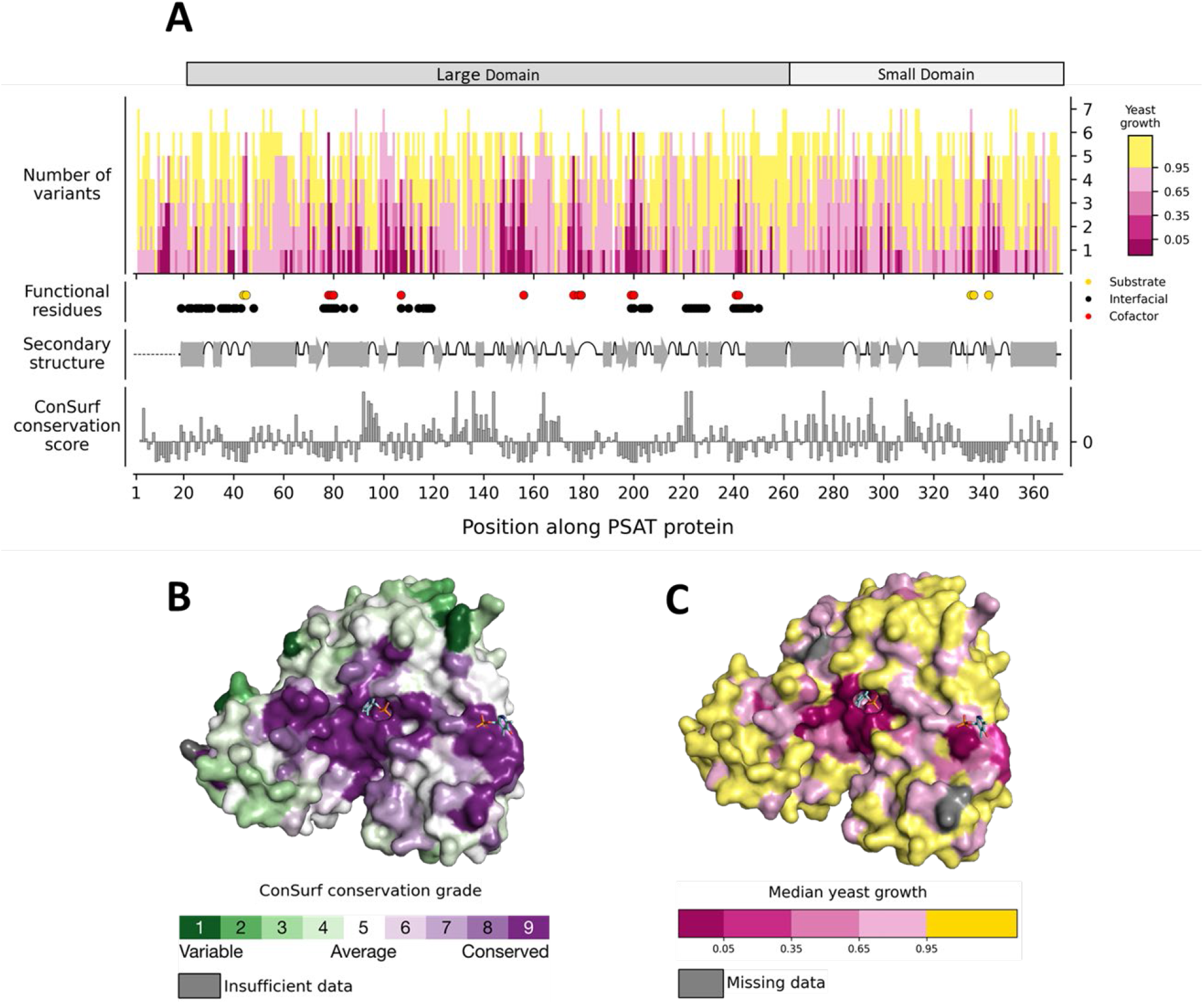
Missense variant effect map across the length of human PSAT compared to structural features and conservation. **A** The top plot depicts the growth of the possible 4-7 substitutions introduced at each amino acid position, ordered from highest (top) to lowest (bottom) growth. Overlayed below are the functional residues of PSAT, secondary structure, and ConSurf evolutionary conservation scores. Subunit domains are indicated above. Residues implicated in substrate binding, formation of the dimer interface, or cofactor binding are shown as circles (righthand legend). Helices and beta-sheets are depicted as gray cylinders and arrows, respectively. Turns shown as upward half-coils. The black dotted line represents secondary structures that were unavailable in the solved crystal structure (PDB: 3e77; L17-L370). More negative ConSurf scores indicate more conserved positions. **B** Subunit surface colored by the ConSurf evolutionary conservation grade and **C** subunit surface colored by median yeast growth estimate per position. Two PLP cofactors are shown in each surface representation (**B,C**) to indicate the other interfacial active site bound to the opposite subunit (not depicted).

The overall structural organization and active site architecture of PSAT are conserved with other members of the class IV family of aminotransferases [42], and phosphoserine aminotransferases from eukaryotic and prokaryotic organisms [55]. To examine our functional scores in the context of evolutionary conservation, we used ConSurf [53] scores (Methods), which are derived from amino acid sequence alignments of homologs. As illustrated in Fig. 4, more conserved (negative ConSurf score) regions localize near the interfacial active sites of the PSAT protein. We expected that highly conserved residues in PSAT would be sensitive to amino acid substitutions. Consistent with this hypothesis, there was a highly significant correlation (Spearman rank correlation=0.47, p < 7.2 × 10^-22^, Additional File 2: Fig. S2A) between the ConSurf score for a residue and the median yeast assay score for substitutions at that residue. As expected, more conserved residues displayed lower median growth. Examining the relationship between the ConSurf scores at a residue and the growth of individual substitutions at that residue further resolved this relationship (Additional File 2: Fig. S2B). Amino acid substitutions that impaired PSAT activity were concentrated at highly conserved residues within PSAT (Fig. 4). However, while few amino acid substitutions at weakly conserved residues exhibited impairment of PSAT function, many substitutions at conserved positions did not (Fig. 4 and Additional File 2: Fig. S2B).

We next considered results of our assay in the context of protein structure. We analyzed a subset of highly conserved functional positions around the active sites, where we see a concentration of residues intolerant to substitutions (Fig. 4). This subset consisted of 12 cofactor binding residues and 5 substrate binding residues. Consistent with the expectation that these sites would be sensitive to amino acid substitutions, we observed that the median growth of variants at these positions was only 7.3%, significantly, and very substantially, lower than the global median of 93.6% (p< 1×10^-5^, one sided permutation test).

We next examined the functional impact of variants at each of these positions individually. In the crystal structure of PSAT complexed with PLP (PDB: 3e77), 12 residues directly interact with the functional groups of the cofactor (Fig. 5A and Fig. 5B) either in pyridoxal ring binding/ coordination or in phosphate group binding. We expected that amino acid substitutions at residues involved in pyridoxal ring binding or coordination would be sensitive to amino acid substitutions (Fig. 5A), as this functional group is directly involved in catalysis [42,56,57]. In fact, all tested substitutions at these six residues (K200, D176, T156, W107, S178, and S179) were functionally impaired. The majority of these, (19/29), including all SNV-accessible missense variants at the catalytic lysine (K200) were amorphic (Fig. 5C). Next, we examined the six residues that participate in phosphate group binding, which may not participate directly in catalysis and whose functional role is less well understood (Fig. 5B). The phosphate group may act as an anchor point to the protein [58], or even directly interact with nearby substrates [57]. At three of these positions (G78, G79, and Q199) all variants were functionally impaired and had amorphic median growth scores (Fig. 5C). Functionally impaired variants were also observed at C80, N241*, and T242*(asterisk indicates residues from opposite subunit), although some substitutions were tolerated (Fig. 5C). Interestingly, most substitutions at the C80 residue are not impaired relative to wild type with the only exception being C80R (growth score=80%). It has been hypothesized that C80 contributes to the higher phosphoserine substrate affinity (Km = 5 μM) observed in human PSAT relative to other phosphoserine aminotransferases [55], and possibly the positive charge introduction represented by C80R negatively impacts this binding.

**Fig. 5.**
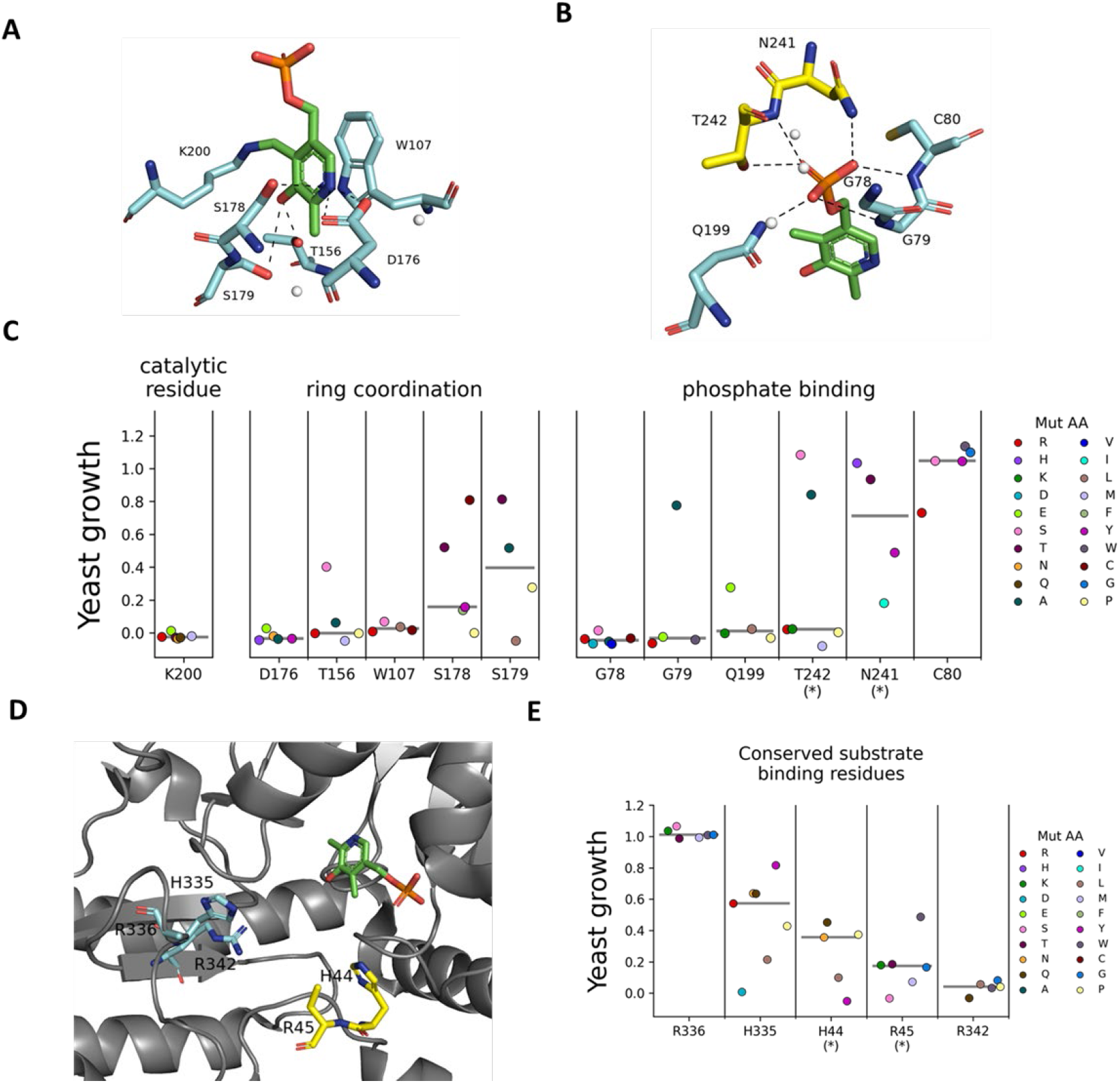
Functional impact of amino acid substitutions at active site residues. Stick representations for residues in one subunit are colored in cyan, residues from the opposite subunit in the homodimer assembly are in yellow, and the PLP cofactor in green in all panels where the crystal structure of human PSAT complexed with PLP (PDB: 3e77) is visualized. **A-B** Organization of PLP cofactor binding residues. Hydrogen bonding is depicted by dashed black lines and nearby water molecules are represented by white spheres. **A** Residues that bind or coordinate PLP’s heterocyclic ring, with the catalytic lysine shown covalently bound to PLP. **B** Residues that bind to the phosphate group of PLP. **C** The distribution of variant growth scores for cofactor binding residues. Each circle is colored according to the amino acid that it is substituted as shown (Mut AA). Asterisks here denote amino acids from the other subunit. The median growth score for each position is shown as a horizontal grey bar. **D** Relative position of highly conserved substrate binding residues to PLP, in the unbound (no substrate) crystal structure of human PSAT. **E** The distribution of functional effects for these conserved substrate binding residues, with the same coloring scheme and annotation as in **C**.

Because the human PSAT crystal structure was solved only without bound substrate, we considered 5 conserved substrate binding residues that were identified in *Escherichia coli* [59], *Bacillus alcalophilus* [60], and *Arabidopsis thaliana* [61] phosphoserine aminotransferases that had crystal structures of phosphoserine or α-methyl-L-glutamate (analog) bound states. Amino acid substitutions at all but one of these sites, corresponding to H335, R336, R342, H44* and R45* (asterisk indicates residues from opposite subunit) in human PSAT, show strong loss of function (Fig. 5D and Fig. 5E). The exception is R336, which displayed activity similar to wild type for all tested substitutions. Thus, R336 may be a conserved residue that does not directly participate in substrate stabilization in PSAT, but does in other orthologs.

The solved human PSAT crystal structure also lacks the first 16 N-terminal amino acids (PDB: 3e77), in which we observe a cluster of sites intolerant to amino acid substitution (residues 11-14) (Fig. 4A). AlphaFold [50,51] predicts that these residues are located in the small domain, proximal to the active site (Additional File 2: Fig. S3). Interestingly, a deletion of 4 N-terminal residues in *Entamoeba histolytica* phosphoserine aminotransferase, corresponding to P12-A15 in human PSAT, yielded a mutant enzyme that had reduced substrate (phosphoserine) affinity, as well as a 10-fold reduction in activity [62]. Additionally, a serine residue involved in analog substrate has been identified in *E. coli* [59], corresponding to P12 in human PSAT. Thus, residues within the first 16 N-terminal residues of human PSAT protein may be involved in substrate binding.

### Mapping functional effects to clinical interpretations

We next compared our results to clinical classifications for all corresponding missense variants in ClinVar. We expected that substitutions derived from known pathogenic variants would produce strong reductions in PSAT activity in our assay, while those derived from known benign variants would have activities comparable to wild type or display only weak reductions in activity. Of the 90 amino acid substitutions derived from variants currently in ClinVar (November 3, 2022, release), three (S179L, A99V, and D100A) are from variants annotated as pathogenic/likely pathogenic, two (I123V and P295R) from variants annotated as benign/likely-benign, and three (V149M, A234S, and R306C) from variants with conflicting interpretations [3]. Since our previous study [39], while the small number of missense alleles with definitive clinical significance calls remained largely unchanged, the number of missense VUS increased nearly 12-fold (from 7 to 82). Consistent with expectation and our previous results [39], the pathogenic/likely pathogenic substitutions all resulted in substantial functional impairment in our assay, with normalized growth values of 82% or below. Similarly, the benign/likely-benign substitutions demonstrated little impairment, with normalized growth values above 91% (Fig. 6).

**Fig. 6.**
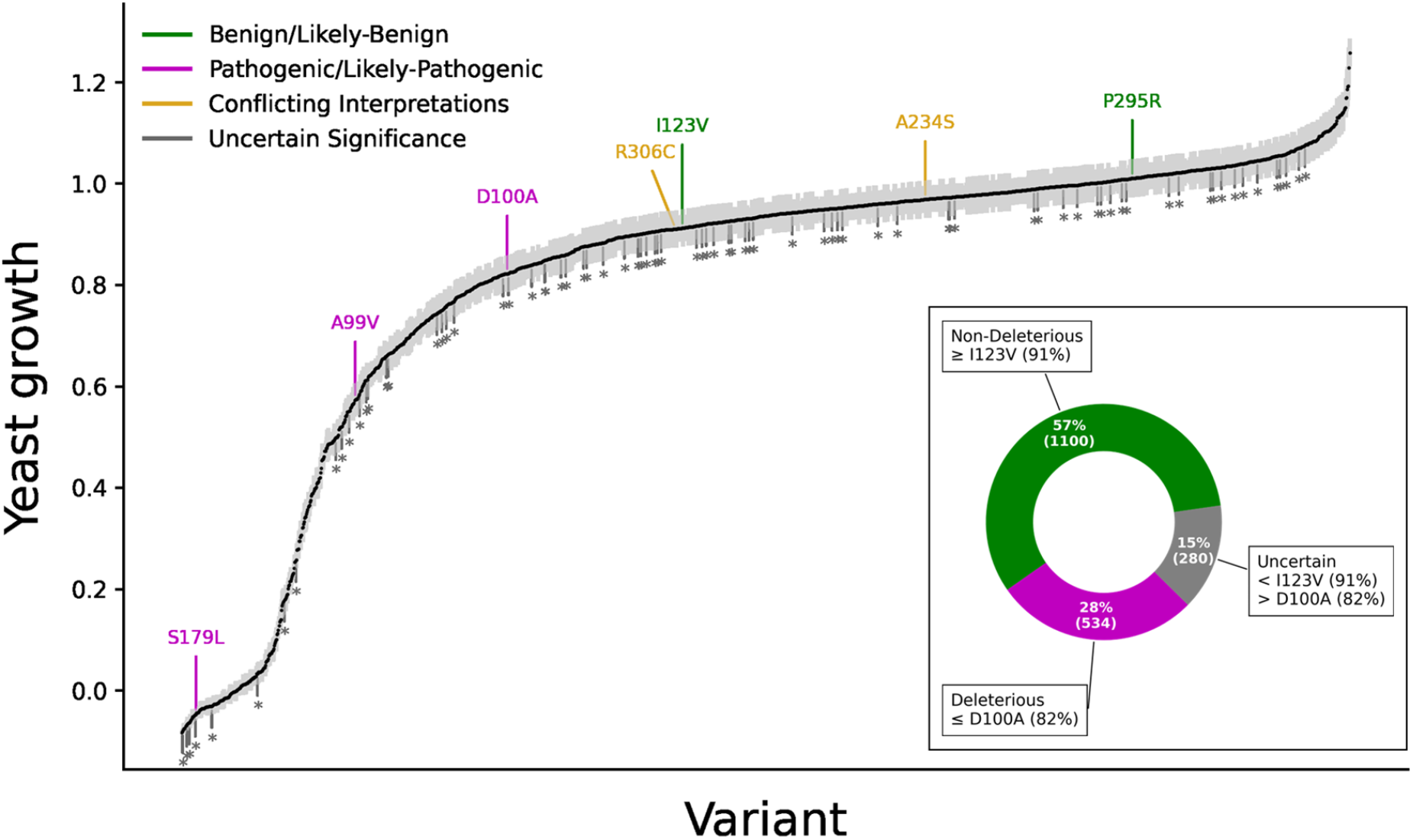
Mutational scan of 1,914 PSAT1 unique SNV-accessible missense amino acid substitutions. Rank ordered (lowest to highest) normalized variant growth. Black dots represent the mean normalized growth estimate for each substitution, with light grey bars indicating standard errors. All current ClinVar annotations associated with tested substitutions are labelled above their growth estimate as benign/likely-benign (green), pathogenic/likely-pathogenic (magenta), conflicting interpretations (yellow), or as an asterisk below their growth estimate if they are of uncertain significance (dark grey). The boxed panel depicts the stratification of normalized growth estimates, based on thresholds derived from ClinVar clinical annotation, into functionally unimpaired (green), indeterminate (gray), or functionally impaired (magenta) ranges.

Data from validated functional assays are potentially valuable as supporting evidence for variant annotation according to the ACMG guidelines [36]. A challenge for rare diseases, such as serine deficiency disorders, is that the limited number of well characterized pathogenic and benign variants precludes the use of current methods, such as odds of pathogenicity [63], for deriving formal measures of classification confidence. As an alternative approach for providing guidance for the use of our data in clinical interpretation, we used a simple thresholding approach, similar that that used in our previous study [39]. Guided by the assay scores of amino acid substitutions with definitive clinical significance calls (Fig. 6), we classified as deleterious any substitutions with ≤82% growth in our assay, which is less than or equal to the scores of substitutions derived from known pathogenic variants (Fig. 6). We also defined a non-deleterious range for scores ≥91%, which is greater than or equal to the scores of substitutions derived from known benign variants (Fig. 6). Because increases in enzyme activity above wild type have not been associated with serine deficiency disorders, we included values greater than wild type in the non-deleterious range. Finally, because there are no clinically annotated variants associated with substitutions having scores between 82% and 91%, this range of the assay is of unknown clinical significance, and we have labeled this range uncertain. Within these ranges of our assay, 534 (28%) of all amino acid substitutions are deleterious, 1,100 (57%) are non-deleterious, and 280 (15%) are uncertain (Fig. 6). Of the 80 ClinVar missense variants currently annotated as VUS that were tested in our assay, 22 result in substitutions that fall in our deleterious range, 41 in our non-deleterious range and 17 in the uncertain range. Thus, our dataset provides functional information for a large number of variants that lack clinical interpretation, including 77% of the missense VUS currently listed in ClinVar.

### Experimental models of homozygous and compound heterozygous genotypes

Pathogenic variants in *PSAT1* cause disease that ranges from severe (Neu-Laxova syndrome 2, NLS2) to milder forms (PSAT deficiency, PSATD). These are collectively referred to as NLS2/PSATD, or individually when clinical severity is discussed specifically. Because NLS2/PSATD is an autosomal recessive disease, clinical manifestation depends on the enzymatic function conferred by the combination of *PSAT1* alleles in the patient’s genome. The ability to generate stable diploid yeast cells by mating haploids allows us to model diploid human *PSAT1* genotypes in yeast and quantitively assess the function of allele pairs. As in our previous study [39], we used our growth assay to functionally assess pairwise combinations of protein-coding variants across all reported unique patient (and carrier parent) genotypes available at the time (Additional File 1: Table S4). Since our previous study [39], six additional reports [28–30,64–66] have added descriptions of 20 new NLS2/PSATD patients and twelve unique genotypes to the disease literature (Additional File 1: Table S4). To experimentally measure the functional impact of these allele combinations in our yeast assay, we constructed diploid strains harboring *yPSAT1* allele combinations encoding the same pair of PSAT1 amino acid sequences as the human genotypes. We then assayed the growth of these strains relative to homozygous wild type (*yPSAT1*) and null (*ser1Δ0*) on minimal medium lacking serine. Previously constructued *yPSAT1* diploid strains encoding homozygous A99V or S43R, and the compound heterzygote A99V / S179L [39], were also included for comparision. Although the growth of strains modelling some patient genotypes was close to that of strains modelling some carrier genotypes, all modelled patient genotypes (Additional File 1: Table S5) displayed reduced growth relative to their respective carrier parents and the homozygous wildtype (Fig. 7B). Consistent with our previous study [39], we also observed good agreement between the degree of *PSAT1* functional impairment in our diploid assay and disease severity (Fig. 7B). Diploids corresponding to genotypes from NLS2 patients had normalized growth from 0% to 77%, and diploids corresponding to genotypes from PSATD patients had a higher normalized growth range of 70% to 88%.

**Fig. 7.**
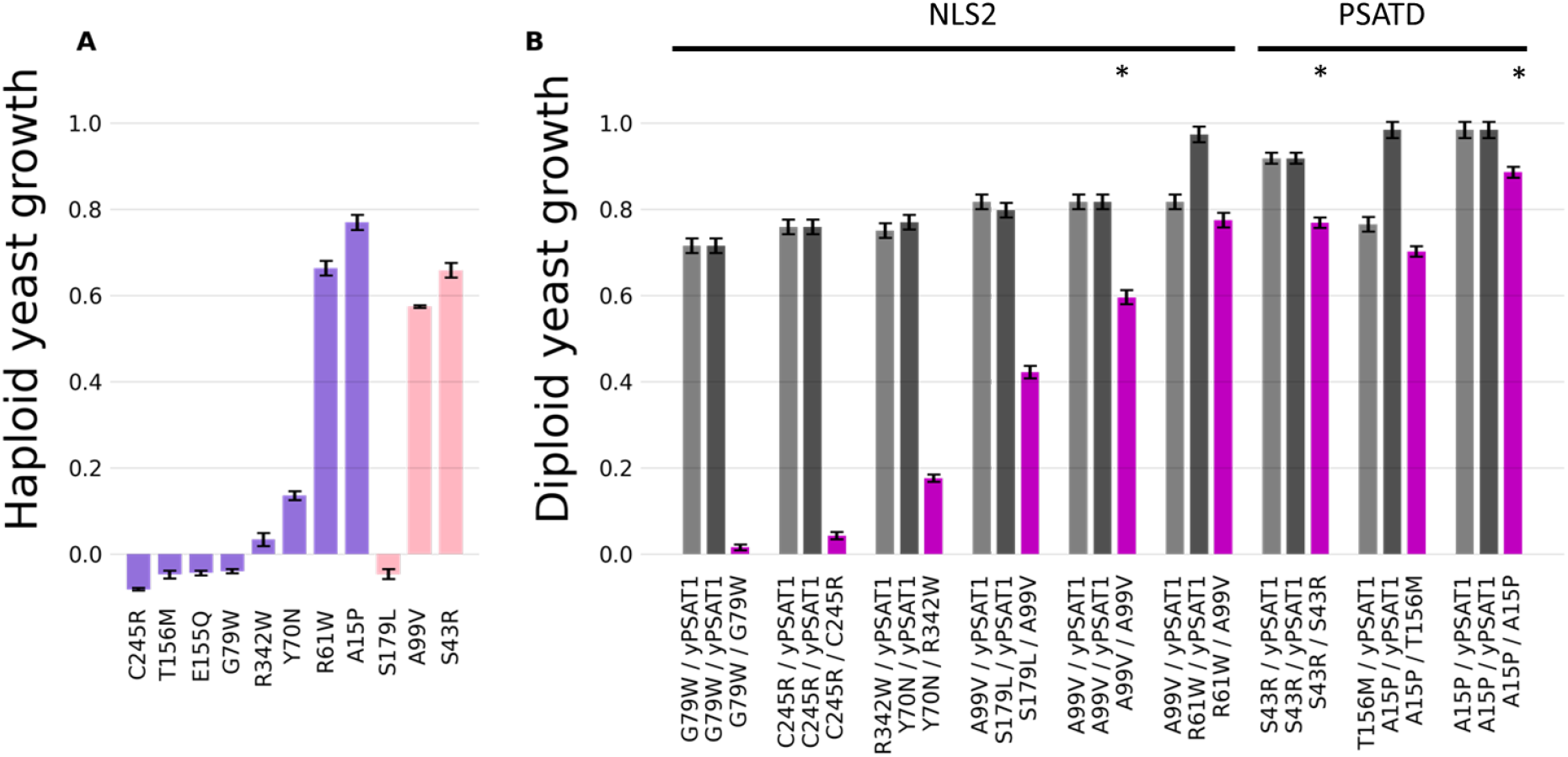
Modeling PSAT activity in patient genotypes. Both bar plots show the mean normalized growth estimate and standard errors. **A** Haploid growth estimates for novel (purple) [28–30,64] and established [25,26] (pink) missense alleles in patient amino acid substitutions. **B** Diploid yeast models of patient trios. Carrier parents are shown in light and dark grey, and the offspring in magenta. Recurrent carrier genotypes are repeated for ease of reference relative to the offspring. The clinical severity (NLS2 or PSATD) associated with patient genotypes is indicated. Asterisks denote offspring genotypes that have been reported in multiple unrelated families.

Our assay results are also consistent with clinical stratification within the NLS2 and PSATD patient groups. For PSATD, the A15P / A15P homozygous genotype is the least impaired in our assay (88% growth) and is also the mildest form of PSATD reported to date. A recent report [30] described two unrelated patients that, other than congenital or childhood ichthyosis, were developmentally normal and displayed neuropathy onset at ages 16 and 17. For NLS2, the two genotypes with the lowest growth in our assay (G79W / G79W and C245R / C245R) correspond to patients displaying severe NLS2 [28]. Three of the remaining NLS2 genotypes modelled here (Y70N / R342W, A99V / S179L, A99V / R61W) display higher growth in our assay and correspond to patients described as having moderate NLS expression [25,28,64]. Among these, Y70N / R342W was the most impaired in our assay (18% growth), and the patient had a postnatal survival of 8 weeks [64]. In contrast, the individual harboring R61W / A99V (77% growth, least impaired) had the longest NLS2-associated postnatal survival described to date (4 months) [28]. Patients homozygous for the A99V genotype (60% growth) exhibit variable postnatal survival (ranging from 1 day to 9 weeks) [25,28,32]. Together, these results highlight the potential value of our model organism assay for variant interpretation in the context of diploid *PSAT1* genotypes.

### Predicting biallelic effects from individual allele measurements

Experimentally assessing the level of PSAT activity associated with all allele-pairs would require construction and assays of 1.83 million diploid yeast strains. As a more labor and cost-effective alternative, we evaluated whether a relationship exists between individual haploid estimates and their resulting diploid growth in combination. For homozygous diploids, a linear relationship was seen between haploid and diploid growth estimates (Fig. 7A and Fig. 7B and Additional File 2: Fig. S4; R^2^=0.98). To extend this analysis to heterozygous genotypes, we fit a model to our experimental diploid dataset of both patient and carrier genotypes (n = 23), to predict diploid growth as a linear combination of the allele with the higher and the allele with the lower haploid growth values (Fig. 8A). This model assumed a uniform degree of dominance of the higher growing allele over the lower growing allele and explained 97% of variance in the growth of the diploids. To compare how this model performed relative to alternatives with simpler assumptions, we also generated models that assumed either complete dominance of the higher-growing allele or an equal contribution of both alleles (Fig. 8B and Fig. 8C). Our full model performed significantly better than either of the alternative models (Anova, Df=1; p=<5.4×10^-6^ and p<2.3×10^-6^, respectively). Leave one out cross validation of the pairwise model displayed an excellent ability to predict diploid growth from haploid measurements (RMSE = 0.0698 and MAE=0.0566).

**Fig. 8.**
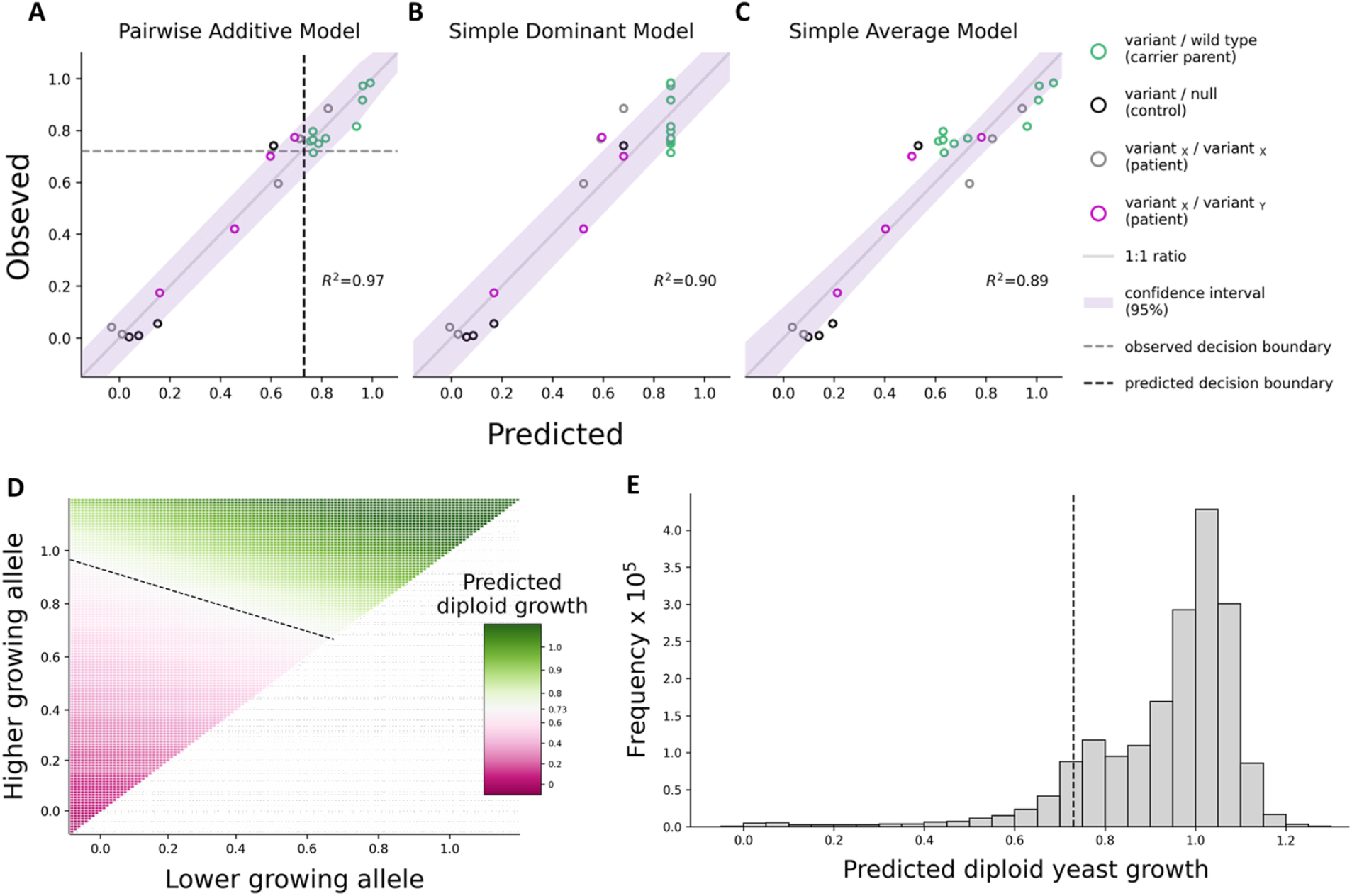
Experimental and predicted biallelic growth values as clinical classifiers. Observed versus fitted values shown for three linear regression models (pairwise additive, **A**; simple dominant, **B**; and simple average, **C**) predicting diploid growth (allele pairs) as a function of haploid growth (single alleles). Each circle represents a unique biallelic combination. Patient, carrier, and control strains are labeled as indicated. A diagonal line of perfect correspondence (1:1 ratio) and the coefficient of determination (R^2^) for each model are included for ease of comparison. Decision boundaries for observed (72%) versus predicted (73%) diploid growth as a binary classifier for identifying patient genotypes modeled in our diploid assay are shown as horizontal and vertical dotted grey and black lines in **A**, respectively. Confidence intervals (95%) for each model are shaded in purple. **D** Heatmap of the predicted diploid growth value for all possible pairwise combinations of haploid allele measurements. **E** Frequency of predicted diploid growth values (in 5% bins) for all ^~^1.8M combinations of alleles. The decision boundary of 73% is shown as a black dotted line in **D** and **E**.

We next compared the results of applying logistic regression to experimental and predicted diploid growth values to produce binary classifiers capable of distinguishing the low growing genotypes of NLS2/PSATD patients (n=9) from the high growing carrier parent genotypes in our trio models (n=10). The threshold growth values at the decision boundary (p=0.5) were similar for both approaches: 72% for experimental and 73% for predicted diploid growth. Interestingly, the decision boundary using predicted values performed better than that of the experimental values, misclassifying only one patient genotype (A15P / A15P) versus four for the model using experimental values (Fig. 8A). It is likely the haploid estimates used to make predictions on diploid growth were more accurate than the experimental diploid estimates as they were generated from a larger number of experimental replicates. These results suggest that diploid growth values predicted from haploid measurements perform well in clinical classification of human *PSAT1* genotypes.

On this basis, we computed the predicted diploid growth from all unique pairwise combinations (^~^1.8 million) of haploid estimates from 1,914 *yPSAT1* missense alleles, wild type (*yPSAT1*), and null (*ser1Δ0*) using the pairwise-additive model (Fig. 8A). We then classified whether the resulting values fell above or below the disease classification boundary of 73% growth. A large majority (89%) of genotypes fell above the decision boundary threshold, suggesting these genotypes would not lead to NLS2/PSATD (Fig. 8D and Fig. 8E). However, over 200,000 *yPSAT1* biallelic combinations displayed predicted diploid growth below the classification threshold, consistent with these genotypes resulting in disease (Fig. 8E).

## Discussion

Here, we present a comprehensive functional analysis of amino acid substitutions associated with SNVs in human PSAT. Leveraging such data for clinical interpretation requires an understanding of the relationship between enzyme function and clinical presentation. Our dataset shows clear stratification of existing clinical variants, with benign variants exhibiting little to no functional impairment, and pathogenic variants exhibiting substantial loss of activity. On this basis, we were able to use the small number of variants with existing clinical interpretation to provide evidence for the likely clinical impact of the large number of variants in similar ranges of the assay. As additional clinical information becomes available through identification of new patients and additional research into the natural history of the disease, the ability to make clinical inferences using our dataset will improve. In the three years since our original study [39], additional patients were described in the disease literature. Not only was there good agreement between the results of our assay and disease severity in these patients, but this additional clinical information also allowed us to extend the range of our assay associated with clinical outcomes.

Given that the therapeutic potential for this medically actionable disorder is highest before patients become symptomatic, clinical sequencing coupled to informative variant interpretation can be a powerful diagnostic tool. Sequencing based diagnostics are becoming more widely applied in newborns, prenatal care [67], and as a part of clinical research efforts such as the Undiagnosed Disease Program [68,69] or Deciphering Developmental Disorders [70]. The most recent AMCG guidelines [71] recommend exome or genome sequencing as a first or second tier option for patients displaying early neonatal (<1 year) congenital abnormalities and for patients exhibiting developmental delay or intellectual disability before the age of 18. In contrast to frameshift and stop-gain mutations, when novel missense variations are identified, even in disease-causing genes, predicting the likely effect on human health is challenging. Relating genotype to phenotype is further complicated by the fact that autosomal recessive diseases are a function of the allele combination in the context of a diploid genome. We can experimentally assay allele combinations in our diploid assay and successfully stratify clinical genotypes.

Furthermore, we were able to predict the effects of allele combinations using a computational model that utilizes combinations of haploid measurements. As a result, we were able to leverage 1,914 yeast measurements to make predictions on 1.8 million potential human genotypes. Given ongoing genome sequencing efforts like the All of US research program and the UK biobank [72], studies that provide information both about individual alleles and the functional impact of allele pairs will play an increasingly important role in realizing the promise of precision medicine.

## Conclusion

The strong agreement between experimentally measured results from our assay and known features of the protein structure together with (albeit limited) clinical data support its use as a well-validated assay of human PSAT function. The corresponding dataset provides meaningful functional information for a large proportion (88%) of all unique SNV-accessible missense substitutions across the length of the protein. Furthermore, our computational model extends the use of the dataset to predict the functional impact of ^~^1.8 million allele combinations corresponding to potential human genotypes. As such, this approach leverages a relatively small amount of clinical data to classify large numbers of variants making it especially powerful for rare diseases.

## Supplementary information

**Additional File 1: Table S1.** *S. cerevisiae* strains used in the haploid assay. **Table S2.** *S. cerevisiae* strains used in the diploid assay. **Table S3.** Normalized *yPSAT1* haploid growth scores and standard error estimates. **Table S4.** Disease literature review of patients diagnosed with PSAT1-related serine biosynthesis defects. **Table S5.** Normalized *yPSAT1* diploid growth scores and standard error estimates. **Table S6.**Human PSAT structure, conservation, and median haploid yeast growth data per amino acid position.

**Additional File 2:** Further description of the MinIon sequencing pipeline. **Fig S1.** Agreement between yPSAT1 variant codons identified by Oxford Nanopore versus Illumina sequencing. **Fig S2.** Comparison of haploid yeast growth scores and ConSurf conservation scores for each corresponding PSAT amino acid position **Fig S3.** Predicted subunit structure for human PSAT.**Fig. S4.** Comparison of experimentally measured haploid and diploid growth scores for PSAT variants that are in homozygous patient (NLS2 or PSATD) genotypes.

**Additional File 3:** Growth normalization script.

**Additional File 4:**MinIon variant calling script.

## Abbreviations

3-PHP: 3-phosphohydroxypyruvate
ACMG: American College of Medical Genetics
cDNA: Protein coding sequence
Df: Degree of freedom
DNA: deoxyribonucleic acid
gnomAD: Genome Aggregation Database
IEM: Inborn errors of metabolism
MAE: Mean absolute error
MATa: Yeast mating type a
MATα: Yeast mating type alpha
NA: Not applicable
NLS: Neu-Laxova syndrome
NLS2: Neu-Laxova syndrome 2
ORF: Open reading frame
PCR: Polymerase chain reaction
PGDH: Phosphoglycerate dehydrogenase
PLP: Pyridoxal Phosphate
PSAT: Phosphoserine aminotransferase
PSATD: Phosphoserine aminotransferase deficiency
PSP: Phosphoserine phosphatase
R^2^: Coefficient of determination
RMSE: Root mean squared error
SD: Minimal medium
SNV: Single nucleotide variant
VUS: Variants of uncertain significance
YPD: Rich medium
yPSAT1: yeast codon-optimized version of the human phosphoserine aminotransferase isoform 1 cDNA sequence

## Declarations

### Ethics approval and consent to participate

Not applicable. No human or animal subjects were used in this work.

### Availability of data and materials

Provided upon manuscript acceptance.

### Authors’ contributions

AMD conceived the project. MJX constructed and assayed the variant library. MJX and MST performed the DNA sequencing. AMD supervised all experimental work. GAC, MT, and KO developed computational pipelines. JNK supervised software development. GAC and MJX performed all data analysis with supervision by AMD. MJX performed all protein structure analysis with supervision from RNM. GAC and MJX performed all comparisons to published clinical data with oversight by AWE-H. MJX, GAC, and AMD wrote the manuscript and incorporated comments from AWE-H, RNM and JNK. All authors approved the manuscript.

## Acknowledgements

We thank Douglas Fowler, Maitreya Dunham, Russell Lo, and Fowzan Alkuraya for helpful discussions.

