## Additional File 2 for "Predicting the functional effect of compound heterozygous genotypes from large scale variant effect maps"

**MinIon Sequencing Pipeline**

A pooled approach utilizing Oxford Nanopore MinIon sequencing was used to identify the sequence variant present in each *YPSAT1* transformant. First, the 96-well variant plates (1 transformant per well) were collapsed in sets of 12, so that well A1 on each collapsed plate combined the variants in well A1 in each of 12 variant plates. Variant plates were ordered so that each of the 12 target codons present in each collapsed well were unique. Therefore, for each collapsed well, we know the 12 target codons present and we also know the set of potential variant sequences introduced at each codon by Twist Biosciences. By sequencing pooled DNA in each well of each collapsed plate and identifying the most common variant sequence at each target codon (among the set of potential variants), we can determine the identity of the variant in each well of the variant plates, i.e. in each transformant.

After collapsing the plates, fragments encompassing the *SER1* promoter, *yPSAT1* ORF, and *SER1* terminator were amplified from the pooled genomic DNA in each collapsed well using 16 or 18 cycles of PCR with dual barcoded primers (identifying each well). ONP adapters were added on via ligation. Sequencing was performed on an Oxford Nanopore MinION Mk1B with a Flongle adaptor & flowcell with 1 collapsed plate per run.

The reads from each sequencing run were demultiplexed to the pool level using MiniBar [1] with the parameters “-l 150 -F -S -M 1 -p .75”. Then, for each pool (collapsed well), reads were aligned to the *yPSAT1* reference sequence in R (R Core Team (2022). R: A language and environment for statistical computing. R Foundation for Statistical Computing, Vienna, Austria. URL https://www.R-project.org/) using the pairwiseAlignment function of the Biostrings package (Pagès H, Aboyoun P, Gentleman R, DebRoy S (2022). Biostrings: Efficient manipulation of biological strings. R package version 2.64.1, https://bioconductor.org/packages/Biostrings) with parameters gapOpening=10 and gapExtension=2. For each read, alignments were carried out in both orientations and the highest scoring alignment was kept. After alignment, at each target codon in the pool, the most frequent variant codon was identified (candidate variant) among the set of potential variant sequences introduced at that codon by Twist Biosciences. The second most frequent variant was also identified. Because we know which unique codon was targeted in each of the 12 transformant strains making up each pool, this gives us a candidate variant for each transformant.

The observed frequency of each candidate variant codon was then compared to the frequency expected under a variant-specific error model. To generate these models, for each pool we recorded the observed frequency of all possible Twist variants at codons NOT among the target set of 12 target codons in that pool. As these variants can only have been produced by sequencing errors, this gives us an error frequency estimate for each potential variant codon. On this basis, provisional variant calls were accepted if their observed frequency was >3.3 times their error frequency. In addition, candidates were rejected when the second most frequent variant was both enriched (>=10-fold) relative to its error frequency and was observed at >30% of the frequency of the candidate variant. Finally, candidates were rejected if they were supported less than 15 reads, or if any missense or nonsense secondary mutations were present in the *YPSAT1* sequence. The script carrying out the MinIon sequence processing steps for each pool (collapsed well) is provided as Additional File 4. At the end of this process, for each transformant, we reported either a high confidence call for the variant present in that transformant, or an NA call resulting in that transformant being removed from analysis.

**Supplementary Figures**

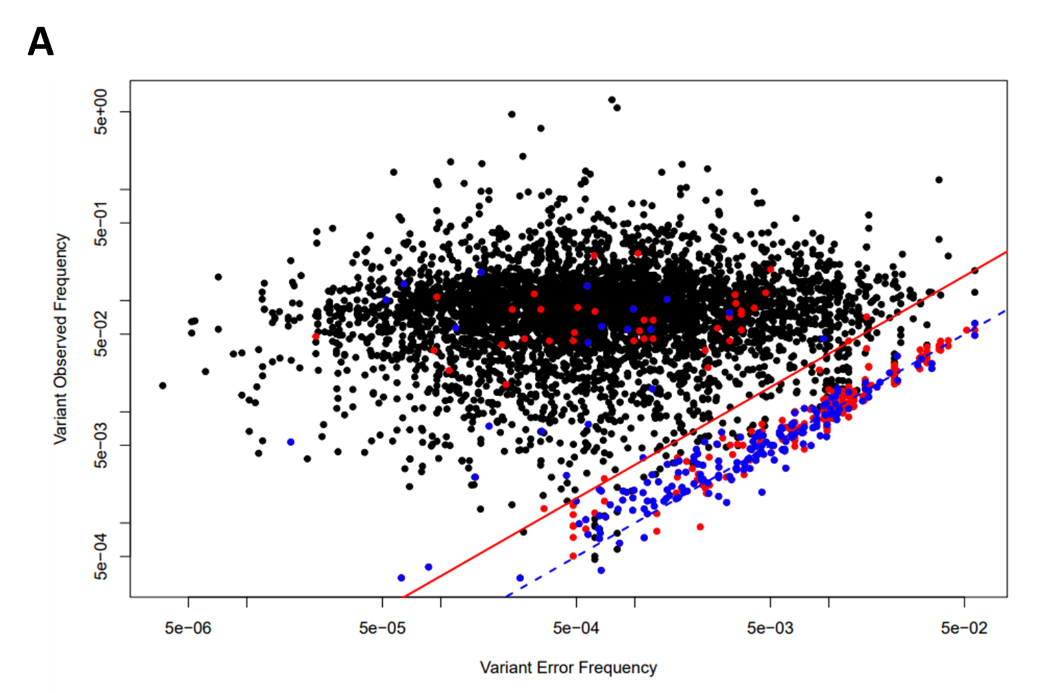

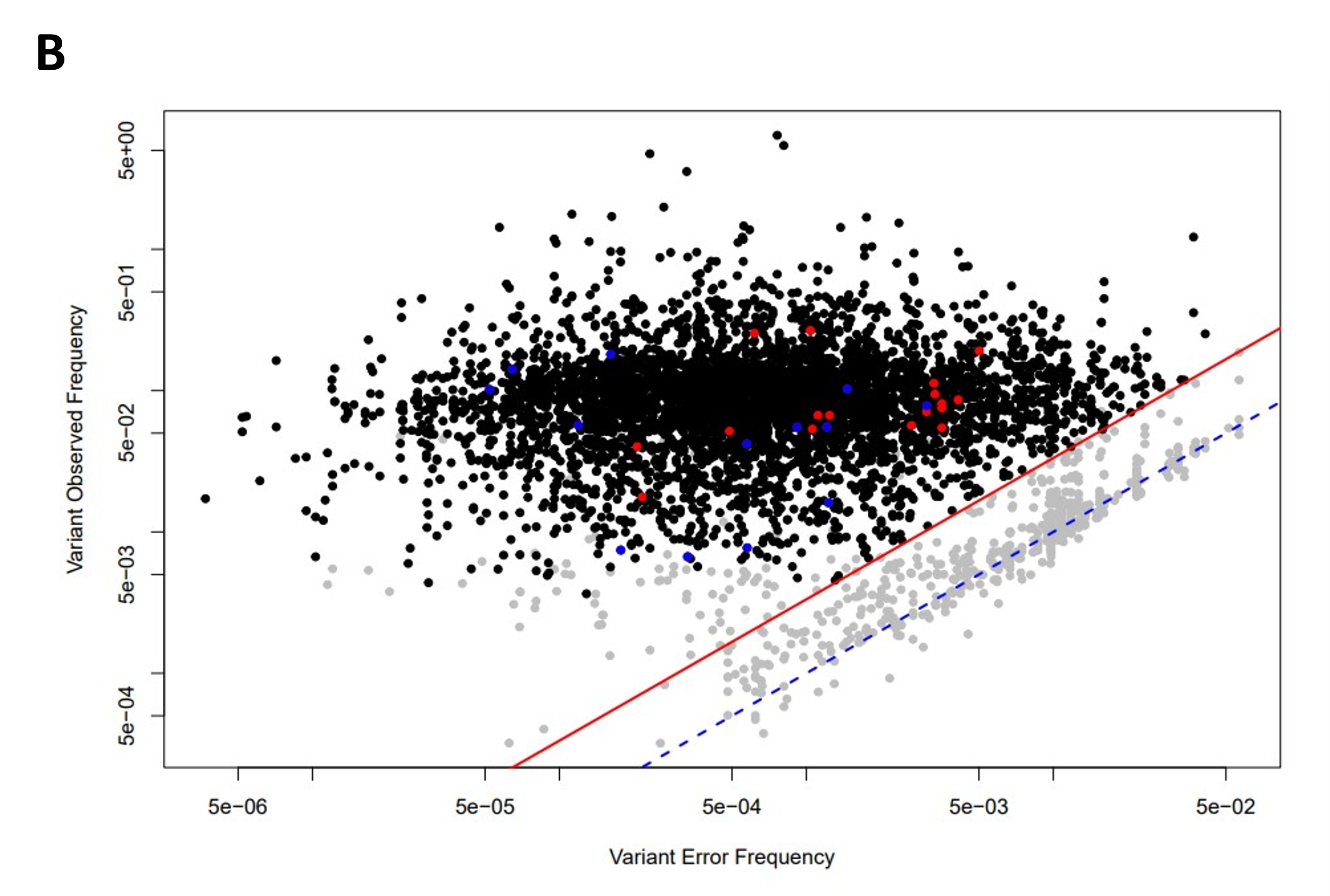

**Fig S1**. Agreement between yPSAT1 variant codons identified by Oxford Nanopore versus Illumina sequencing. For each variant call, the observed ratio of Oxford Nanopore variant to reference reads (expected to be 1/15) is plotted against the empirically calculated error ratio. Blue dotted line indicates 1:1 relationship between observed and error ratios. Quality filters require observed variant frequency to be >3.3 times the error frequency (red line). **A** Variant calls in agreement with Illumina shown in black. Disagreements where Illumina identifies no change at the target codon (i.e. reference sequence) shown in blue. Remaining disagreements shown in red. **B** As in **A** except that variants failing quality control filters are colored in grey.

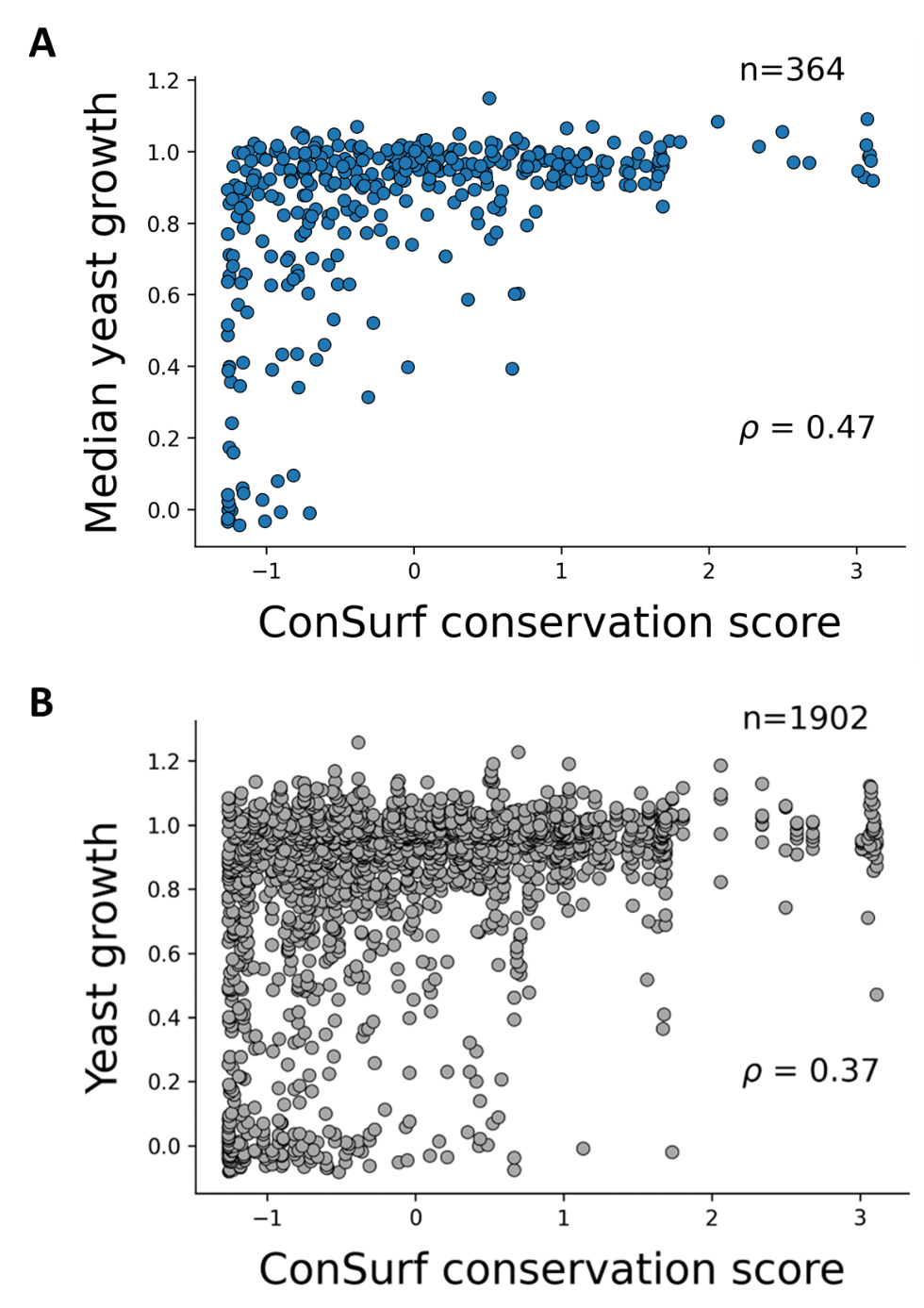

**Fig S2.** Comparison of haploid yeast growth scores and ConSurf [2] conservation scores for each corresponding PSAT amino acid position. More negative conservation scores indicate more conserved scores. **A** Scatter plot of the median yeast substitution score and ConSurf score per amino acid position. **B** Scatter plot of haploid yeast growth scores for each substitution and the ConSurf score for their amino acid position. The corresponding Spearman rank correlation (ρ) is labeled on each plot (p<7.2 x 10^-22^ in **A**, p<1.5 x 10^-62^ in **B**).

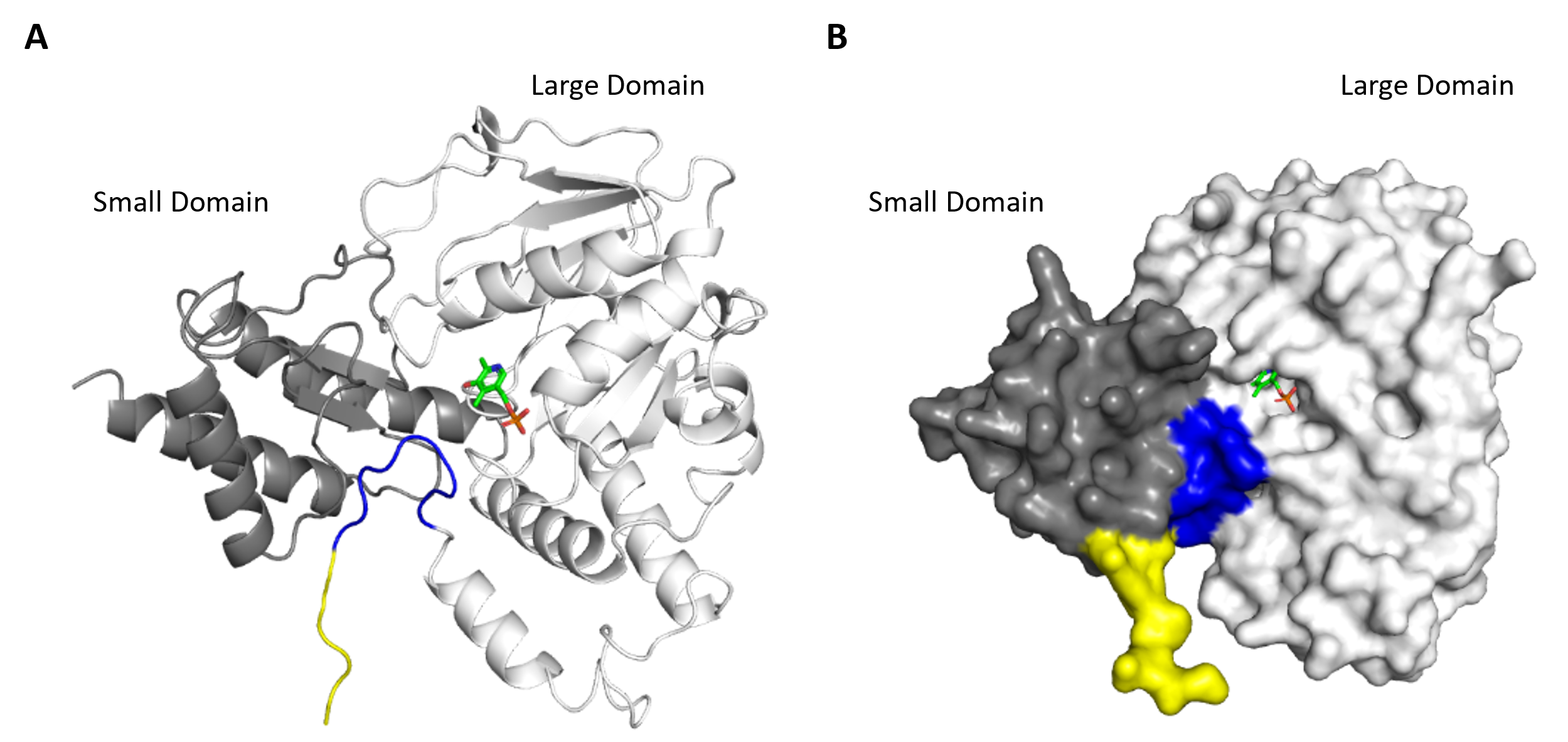

**Fig S3**. Predicted subunit structure for human PSAT. **A** Cartoon and **B** surface representation of the subunit structure for human PSAT (UniProtID=Q9Y617-1, version 2002-01-23) [3], as predicted by AlphaFold (version 2022-11-01) [4,5]. Small and large domains are labeled on both. The PLP cofactor (green) is represented as a stick and was transplanted into the predicted structure using AlphaFill [6], to better visual the location of one of the active sites. Residues with AlphaFold per-residue confidence scores (pLDDT) of very confident ratings (>90) are colored in blue (V8-K16). Residues below this confidence rating of very confident (<90), are shown in yellow (M1-V7).

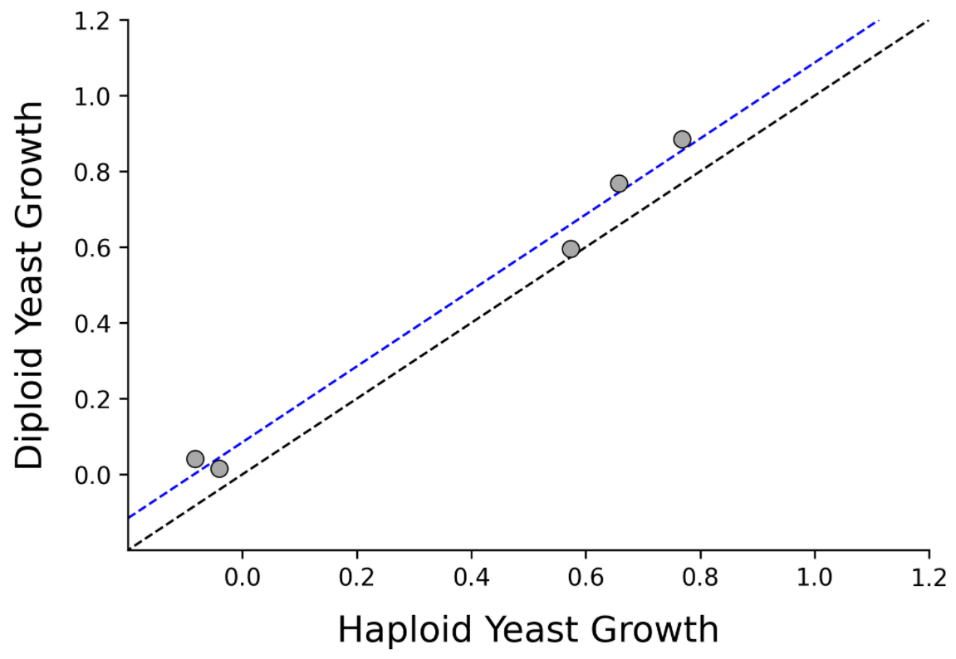

**Fig S4**. Comparison of experimentally measured haploid and diploid growth scores for PSAT variants that are in homozygous patient (NLS2 or PSATD) genotypes. Scatter plot of normalized haploid and diploid growth scores. Haploid scores were scaled relative to wild type *yPSAT1* (normalized growth=1) and null (normalized growth=0). Diploid scores were scaled relative to homozygous wild type *yPSAT1 / yPSAT1* (normalized growth=1) and null / null (normalized growth=0). The black dotted line represents a 1:1 correspondence between the haploid and diploid scores. The blue dotted line indicated the observed correlation (R^2^=0.989) with slope of 1.0 and intercept of 0.09.
